# Akt activator SC79 stimulates antibacterial nitric oxide production from human nasal epithelial cells and increases macrophage phagocytosis *in vitro*

**DOI:** 10.1101/2022.10.31.514559

**Authors:** Robert J. Lee, Nithin D. Adappa, James N. Palmer

**Affiliations:** Department of Otorhinolaryngology, University of Pennsylvania Perelman School of Medicine; Department of Physiology, University of Pennsylvania Perelman School of Medicine

**Keywords:** chronic rhinosinusitis, *Pseudomonas aeruginosa*, *Staphylococcus aureus*, air-liquid interface

## Abstract

**Background:** The role of the Akt serine/threonine kinase family in airway innate immunity is relatively unstudied compared with other pathways. Akt can phosphorylate and activate the endothelial nitric oxide synthase (eNOS) isoform expressed in airway epithelial ciliated cells. NO production by nasal epithelial cells often has antibacterial and antiviral effects. Increasing nasal epithelial NO production may be a useful anti-pathogen strategy for respiratory infections in diseases like chronic rhinosinusitis. We hypothesized that a small molecule Akt activator, SC79, might induce nasal epithelial cell NO production with bactericidal effects.

**Methods:** We tested the antibacterial-stimulatory effects of SC79 in primary nasal epithelial cells isolated from residual surgical material and grown at air-liquid interface. Because macrophages also use NO signaling to enhance phagocytosis, we also tested effects of SC79 in human macrophages differentiated from monocytes obtained from healthy apheresis donors.

**Results:** Live cell imaging of an NO-sensitive fluorescent dye revealed that SC79 induced dose-dependent NO production. Pharmacology and genetic knockdown revealed that this NO production is dependent on eNOS and Akt. The NO released into the airway surface liquid was sufficient to kill both lab and clinical strains of *P. aeruginosa* in a co-culture bacterial killing assay. SC79 enhanced bacterial phagocytosis in a NO-dependent and Akt-dependent manner. No overt toxicity (LDH release) or inflammatory effects (IL8 transcription) were observed in nasal cells or macrophages over 24 hrs.

**Conclusions:** Together, these data suggest that multiple innate immune pathways might be stimulated by SC79 delivered via topical nasal rinse or spray. Activating Akt using SC79 or another compound might have beneficial antipathogen effects in respiratory infections.

## Background

Generation of nitric oxide (NO) is an important airway defense mechanism. NO directly kills or inactivates bacteria and fungi by damaging their cell walls or DNA [1–5]. NO also inhibits replication of influenza, parainfluenza, rhinovirus [6], and SARS-COV1 & 2 viruses [7–10]. NO is generated by the enzymatic conversion of the amino acid L-arginine to L-citrulline by NO synthase (NOS). In healthy nasal ciliated cells, endothelial NOS (eNOS) is the main NOS isoform, constitutively expressed [11–13] and apically localized to the cilia base or within the cilia body itself [14–18]. In endothelial cells, activation of eNOS can occur either by Ca^2+^/calmodulin binding to eNOS or by kinase phosphorylation of eNOS at key residues like serine 1177 (S1177) [19, 20].

We previously showed that bitter taste receptors (taste family 2 receptors, or T2Rs) activate eNOS in airway cilia [21, 22]. T2Rs are G-protein coupled receptors (GPCRs) originally identified on the tongue [23] but also found in airway motile cilia [24–28]. Cilia-localized T2Rs detect bacterial metabolites (e.g., acyl-homoserine lactones [AHLs] [29] and quinolones [27] from *Pseudomonas*) and activate rapid (within minutes) Ca^2+^-dependent eNOS activation to produce NO. This NO both increases cilia beating to clear bacteria and can directly kill bacteria [25, 29, 30]. Macrophage-expressed T2Rs enhance phagocytosis in response to bacterial metabolites through a similar Ca^2+^ and eNOS signaling pathway [31]. Thus, increasing NO generation may be useful in multiple types of cells involved in airway defense.

However, not everyone can generate nasal epithelial NO via T2Rs, and this appears to have detrimental clinical results. Approximately 25% of people are homozygous for the *TAS2R38* AVI polymorphism, which encodes a non-functional version of T2R isoform T2R38, one of the most highly expressed T2R isoforms in sinonasal cilia. AVI/AVI individuals are more susceptible to gram-negative upper respiratory infection [25] and chronic rhinosinusitis (CRS) [32, 33]. AVI/AVI individuals have poorer sinus surgical outcomes than patients homozygous for PAV (functional) T2R38 [34]. This was supported by genome-wide association studies of *TAS2R38* [35, 36] and a study showing AVI/AVI patients have increased sinonasal bacteria [37].

AVI *TAS2R38* may also be detrimental in cystic fibrosis (CF) patients [38, 39]. However, we recently discovered that cells from F508del/F508del CF patients exhibit reduced T2R-stimulated NO production despite similar upstream Ca^2+^ signals [40]. This suggests that impaired NO signaling in CF cells results in reduced T2R responses to bitter bacterial products, including the AHLs and quinolones secreted by *P. aeruginosa*, regardless of *TAS2R38* genotype. We thus hypothesized that “boosting” epithelial NO production by a non-T2R pathway in nasal cells is a potential complementary strategy to treat upper respiratory infections, particularly in *TAS2R38* AVI/AVI patients or in CF-related CRS patients. Activating innate immune responses like NO production to reduce acute or chronic bacterial infections could reduce the pressures for single agent resistance in chronic rhinosinusitis [41–44].

One pathway to activate eNOS, largely unexplored in airway cell physiology, is the Akt kinase pathway. There are three mammalian Akt isoforms (Akt1, 2, and 3), which vary in tissue expression. We previously found that primary nasal cells grown at air liquid interface (ALI) express all three Akt isoforms [45]. Once activated, Akt can phosphorylate multiple downstream targets like other kinases or transcription factors [46, 47], including many proteins associated with cell survival and proliferation [47, 48]. Another target of Akt is eNOS. AKT phosphorylation of eNOS at S1177 can activate eNOS NO production [19, 20]. We hypothesize that the NO production driven by Akt in nasal epithelial cells, like the NO produced during T2R stimulation, may enhance bacterial killing. The data below are an *in vitro* pilot study using a small molecule Akt activator to determine if targeting the Akt pathway might enhance innate immune responses during upper respiratory infections. As a model, we used both primary human nasal epithelial cells as well as primary human monocyte-derived macrophages.

## Materials and methods

### Reagents and Solutions

SC79, SNAP, L-NAME, D-NAME, cPTIO, MK2206, LY294002, U73122, U73343, GSK690693, and colistin sulfate were from Cayman Chemical (Ann Arbor, MI, USA). API-2 was from Bio-Techne/Tocris (Minneapolis, MN, USA). DAF-FM diacetate was from ThermoFisher Scientific (Waltham, MA, USA). Unless specified below, all reagents were from MilliporeSigma (St. Louis, MO, USA). LDH assay kit (catalog no. ab65393) was from Abcam (Cambridge, MA, USA). *P. aeruginosa* flagellin (FLA-PA) was from InvivoGen (San Diego, CA). Stock solutions of hydrophobic compounds were made in DMSO at ≥1000x so that final DMSO concentrations were ≤0.1%. Vehicle control matched the highest DMSO concentration used in an experimental paradigm. Hank’s balanced salt solution (HBSS) was used in all live cell imaging experiments and contained (in mM) 137 NaCl, 5 KCl, 0.4 KH_2_PO_4_, 0.3 Na_2_HPO_4_, 5.5 glucose, 1.8 CaCl_2_, 1.5 MgCl_2_, 20 HEPES, pH 7.4, supplemented with 1x MEM amino acids to provide a roughly physiological level of L-arginine for NO production.

### Cell Culture

Primary human nasal epithelial cells were collected in accordance with The University of Pennsylvania guidelines regarding use of residual clinical material from patients undergoing sinonasal surgery at the University of Pennsylvania with institutional review board approval (#800614) and written informed consent from each patient in accordance with the U.S. Department of Health and Human Services code of federal regulation Title 45 CFR 46.116. Inclusion criteria were patients ≥18 years of age undergoing sinonasal surgery for sinonasal disease (CRS) or other procedures (e.g. trans-nasal approaches to the skull base). Exclusion criteria included history of systemic inheritable disease (e.g., granulomatosis with polyangiitis, cystic fibrosis, systemic immunodeficiencies) or use of antibiotics, oral corticosteroids, or anti-biologics (e.g. Xolair) within one month of surgery. Individuals ≤18 years of age, pregnant women, and cognitively impaired persons were not included. Tissue was transported to the lab in saline on ice and mucosal tissue was immediately removed for cell isolation.

Sinonasal epithelial cells were enzymatically dissociated and grown to confluence in proliferation medium (50% DMEM/Ham’s F-12 plus 50% bronchial epithelial basal media [BEBM] plus Singlequot supplements; Lonza, Walkersville, MD, USA) for 7 days [21]. Confluent cells were dissociated and seeded on Corning (Corning, NY, USA) transwells (0.33 cm^2^, 0.4 µm pore size; transparent) coated with BSA, type I bovine collagen, and human fibronectin. When culture medium was removed from the upper compartment, basolateral media was changed to differentiation medium (1:1 DMEM:BEBM) containing hEGF (0.5 ng/ ml), epinephrine (5 ng/ml), BPE (0.13 mg/ml), hydrocortisone (0.5 ng/ml), insulin (5 ng/ml), triiodothyronine (6.5 ng/ml), and transferrin (0.5 ng/ml), supplemented with 100 U/ml penicillin, 100 g/ml streptomycin, 0.1 nM retinoic acid, and 2% NuSerum (BD Biosciences, San Jose, CA) from the Singlequot supplements as described [21]. Cells were fed basolaterally with differentiation media for ∼21 days prior to use. Primary ALI cultures were genotyped for *TAS2R38* AVI (non-functional) or PAV (functional) polymorphims as described [21]. Differentiation was verified based airway epithelial morphology (formation of motile cilia, goblet cells) and transepithelial electrical resistance, etc..

Primary human M0 macrophages were cultured as described [31] in RPMI2650 media with 10% human serum and 1x cell culture Pen/Strep. De-identified monocytes from healthy apheresis donors were obtained from the University of Pennsylvania Human Immunology core with written informed consent of every participant and institutional review board approval. Cells isolated from 10 different individuals were used. As all samples were de-identified for race, age, sex, etc., samples were used in a blinded fashion. Macrophages were differentiated by adherence culture for 12 days in 8-well chamber slides (CellVis; Mountain View, CA) as described [31]. Our prior studies suggest no differences in T2R responses among macrophages differentiated by adherence alone or by adherence plus M-CSF [31], and thus adherence only was used for these studies.

16HBE cells were cultured in MEM (ThermoFisher Scientific) containing 10% FBS (Sigma) and 1% penicillin/ streptomycin (ThermoFisher Scientific).

### Live cell imaging of Akt activity

The 16HBE cells were originally created by D. Gruenert (UCSF [49]) with CFTR CRISPR modified versions obtained under materials transfer agreement from Cystic Fibrosis Foundation Therapeutics [50]. Cells were plated onto glass 8-well chamber slides (CellVis, Mountain View, CA, USA) and transfected the next day with Lipofectamine 3000 and cerulean/cpV FRET-based sensor AktAR (gift of Jin Zhang via Addgene, Watertown, MA, USA; [51]). Cells were imaged on an inverted microscope (Olympus, Tokyo Japan; 20x 0.8 NA objective) with motorized programmable stage (Prior Scientific, Rockland MA) and standard CFP/YFP emission filters in motorized filter wheels (Lambda LS, Sutter Instruments, Novato California). CFP and YFP emission (both with CFP excitation) was collected every 2 min for 3 hours.

### Live cell imaging of NO production

Primary human ALIs were loaded for 90 min with 10 µM DAF-FM diacetate as previously described [21] in Hank’s Balanced Salt solution (HBSS) buffered to pH 7.4 with 20 mM HEPES. Macrophages and 16HBE cells on glass chambered coverslips were loaded with 5 µM DAF-FM diacetate for 45 min as previously described [27, 31]. DAF-FM was imaged as previously described [21, 52] on a TS100 microscope (Nikon, Tokyo, Japan) with 10x 0.3 NA PlanFluor objective, Retiga R1 CCD camera (Photometrics, Tucson, AZ, USA), standard FITC/GFP filter set (Chroma, Bellows Falls, VT USA), and XCite 110 LED illumination source. Images were acquired with the micromanager variant of ImageJ [53] and analyzed in FIJI [54].

### Immunofluorescence (IF) microscopy

IF was carried out as previously described [21]. ALIs were fixed in 4% formaldehyde for 20 min at room temperature, followed by blocking and permeabilization in Dulbecco’s phosphate buffered saline (DPBS) containing 1% bovine serum albumin (BSA) and 5% normal donkey serum (NDS) for blocking as well as 0.2% saponin and 0.3% triton X-100 for permeabilization for 1 hour at 4°C. Primary antibody incubation (1:100) was at 4°C overnight. Incubation with AlexaFluor (AF)-labeled antibodies (1:1000) was for 2 hours at 4°C. Transwell filters were removed from the plastic ring and mounted with Fluoroshield with DAPI (Abcam; Cambridge, MA USA). Images were taken on an Olympus Fluoview FV1000 confocal system with IX-73 microscope and 60x (1.4 NA) objective and analyzed in FIJI [54] using linear min/max adjustments. Anti-beta-tubulin IV (ab11315; mouse monoclonal) antibody was from Abcam. Donkey-raised, Alexa-Fluor-conjugated secondary antibodies (anti-rabbit AlexaFluor 546 and anti-mouse AlexaFluor 488) were from ThermoFisher Scientific.

### Quantitative PCR

RNA was isolated from ALI cultures or stripped turbinate or polyp epithelium, as described previously [55], and qPCR was performed using TaqMan primers (ThermoFisher Scientific) for IL8 and glyceraldehyde-3-phosphate dehydrogenase in separate reactions, and relative expression was calculated by calculated by means of the 2^ΔΔCt^ method.

### Bacteria culture

*P. aeruginosa* strains PAO-1 (ATCC 15692) and clinical CRS-isolates (Drs. N. Cohen and L. Chandler, Philadelphia VA Medical Center [56]) were grown in Luria broth (Gibco). For anti-bacterial assays, *P. aeruginosa* were grown to OD 0.1, centrifuged, and resuspended in 50% 0.9% saline supplemented with 0.5 mM glucose and 1 mM HEPES, pH 6.5. Nasal ALIs were washed 24 hrs prior to the assaywith antibiotic-free Ham’s F12K media (ThermoFisher Scientific) basolaterally. 30 uL of bacteria saline solution was placed on the apical side of the ALI for 10 min, followed by aspiration of bulk ASL fluid. After 2 hrs at 37 °C, remaining bacteria were collected from the ALI by washing. Bacteria were live-dead stained with SYTO9 (live; green) and propidium iodide (dead; red) with BacLight Bacterial Viability Kit (ThermoFisher Scientific). Green/red fluorescence ratio was quantified in a Spark 10M (Tecan, Mannedorf, Switzerland; 485 nm excitation with 530 nm and 620 nm emission). CFU counting was done as previously described on LB plates with serial dilutions [22, 56].

### Phagocytosis assays

Phagocytosis assay were performed as described [31]. Macrophages were incubated in phenol red-free, low glucose DMEM with heat-killed FITC-labeled *Escherichia coli* at 250 µg/ml (strain K-12; reagents from Vybrant phagocytosis assay kit; ThermoFisher Scientific; Cat # E2861) (ThermoFisher Scientific) ± T2R agonist for 15 min at 37°C. Extracellular FITC was quenched with trypan blue per the manufacturer’s instructions, and fluorescence was recorded on a Spark 10M plate reader (Tecan; 485 nm excitation, 535 nm emission). As phagocytosis is negligible from 4 °C up to room temp [31], we recorded fluorescence from living cells at room temperature immediately after the FITC-*E. coli*. For representative micrograph shown, macrophages on glass were incubated as above, and extracellular FITC was quenched with trypan blue and cells were washed ≥5x in PBS to remove residual extracellular FITC-*E. coli*. Remaining adherent MΦs were fixed in 4% formaldehyde (Electron Microscopy Sciences, Hatfield, PA) for 10 min followed by DAPI staining in mounting media (Fluoroshield with DAPI, Abcam). FITC-E. coli were then imaged using standard FITC filter set (Semrock, Rochester, NY USA) on an inverted Olympus IX-83 microscope with 20x (0.8 NA) objective, XCite 120LEDBoost illumination source, and Hammamatsu (Tokyo, Japan) Orca Flash 4.0 sCMOS camera.

Phagocytosis assays were also carried out similarly using 125 µg/ml pHrodo red-labeled *S. aureus* (strain Wood 46; ThermoFisher Scientific, cat # A10010) [31]. As pHrodo dyes only fluoresce when particles are internalized into low pH endosomes (previously demonstrated in [31]), this assay does not require washing or quenching of the extracellular pHrodo *S. aureus*. Macrophages were incubated with pHrodo-*S. aureus* for 30 min at 37°C as described [31] with excitation at 555 nm and emission at 595 nm measured on the Tecan Spark 10M plate reader. Background measurements were made using wells containing fluorescent *S. aureus* in the absence of macrophages. Representative images were taken as above except using a standard TRITC filter set (Semrock). Images shown for comparison were collected on the same day under identical conditions with identical min/max settings. No non-linear (e.g., gamma) adjustments were made to any images for either display or analysis.

### Data analysis and statistics

Multiple comparisons were analyzed with one-way ANOVA with appropriate posttests: Bonferroni for pre-selected pairwise comparisons, Tukey-Kramer for comparing all values, or Dunnett’s for comparing to control value. Asterisks (* and **) indicate *p* <0.05 and *p* <0.01, respectively; *p* <0.05 was considered statistically significant. All data in bar graphs are shown as mean ± SEM with n derived from biological replicates (separate experiments conducted with different patient cells on different days). Raw unprocessed image data were analyzed in FIJI [54] and resulting numerical data were analyzed in Excel (Microsoft, Redmond, WA, USA) and/or Prism (GraphPad software, La Jolla, CA). All data used to generate bar graphs and traces are available upon request.

## Results

We tested direct Akt activation using small molecule Akt activator SC79. SC79 binds to Akt to promote a conformational change allowing Akt phosphorylation by upstream kinases [57–61]. SC79 activation of Akt has beneficial effects in animal models of neuronal [59, 60] and hepatic disease [61]. In A549 lung cancer cells, 16HBE immortalized bronchial epithelial cells, and primary nasal epithelial cells, we previously found that SC79 acutely increased Akt phosphorylation at S473 [45]. The resulting Akt activation caused translocation of the transcription factor Nrf-2 to the nucleus, reducing NFκB-driven IL-8 transcription during co-stimulation with TNFα or toll like receptor 5 (TLR5) agonist flagellin (**Figure 1a**) [45]. We also observed that SC-79 increased eNOS phosphorylation at S1177 and stimulated NO production in A549 cells (**Figure 1b**) [45]. However, the consequence of this NO production were not determined, including whether the NO produced diffuses into the airway surface liquid and whether it is of sufficient magnitude to kill bacteria *in vitro*. In non-CF 16HBE cells containing Wt CFTR (parental cell line) or CRISPR modified to contain F508del or G542X CFTR [50], we visualized SC79 Akt activation using ratiometric Akt biosensor AktAR [51] (**Figure 1c-d**). We observed similar levels of Akt activation with SC79 in all three cell types (**Figure 1e**). While some have reported altered receptor-dependnet Akt signaling in CF cells [62–64], this does not appear to occur with AktAR. We thus hypothesized that SC79 could be used to drive NO production in both non-CF and CF cells. Bitter T2R agonists activate NO production by a Ca^2+^-dependent, Akt-independent pathway [40]. T2R-activation of NO, measured by fluorescent reactive nitrogen species (RNS) indicator DAF-FM, is reduced in CF 16HBE cells (**Figure 1f** and [40]). However, when we stimulated non-CF and CF 16HBEs with SC79 (either 1 or 10 µg/ml), we saw similar levels of NO production among the cells (**Figure 1g**). This validated our hypothesis in isogenic immortalized cells, but further validation required data from primary cells.

**Fig. 1.**
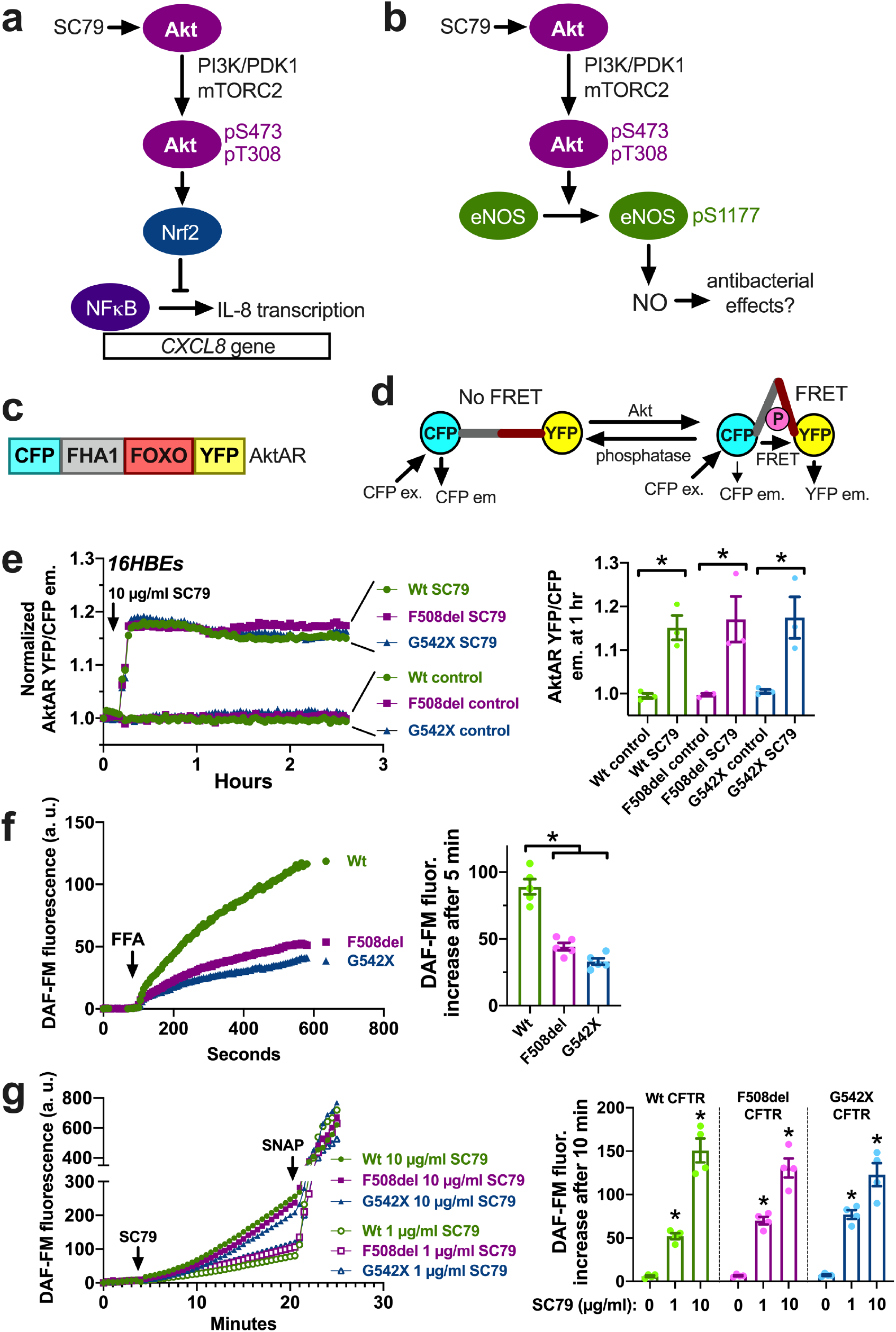
Small molecule Akt activator SC79 increases NO production in 16HBE air-liquid interface cultures (ALIs). **a** SC79 binds to Akt and promotes its phosphorylation and activation downstream of PI3 kinase (PI3K) and phosphoinositide-dependent protein kinase-1 (PDK-1) as well as mTOR Complex 2 (mTORC-2) at S473 and pT308. We previously showed that SC79 causes Akt-dependent increase in Nrf-2 levels in the nucleus that suppress NFκB transcription of IL-8 [45]. **b** We also showed that SC79 induced Akt-dependent phosphorylation of endothelial nitric oxide synthase (eNOS) at pS1177 that stimulates nitric oxide (NO) production. We hypothesized that this may have antibacterial effects in nasal epithelial cells. **c** The AktAR biosensor contains cyan fluorescent protein (CFP) variant cerulean and YFP-variant circularly permutated (cp)Venus with an E172 mutation. These proteins flank a phosphorylated amino acid binding domain created from forkhead-associated domain (FHA1) and and the Akt substrate sequence surrounding Thr-24 of FOXO1. **d** Phosphorylation by Akt causes a conformational change that brings the CFP and YFP closer together increasing the YFP emission and decreasing the CFP emission when CFP is excited. **e** Left, representative traces showing AktAR fluorescence changes in 16HBE cells with Wt, G542X, and F508del CFTR. Right, bar graph showing change in YFP/CFP emission ratio 1 hr after stimulation with SC79; *p<0.05 vs isogenic control by one-way ANOVA with Bonferroni posttest. No differences were observed between baseline or stimulated YFP/CFP ratios among the various CFTR genotypes by one-way ANOVA with Bonferroni posttest. Bar graph shows data points from n = 3 independent experiments using cells at different passages. **f** Left, representative DAF-FM fluorescence trace during stimulation with 100 µM flufenamic acid (FFA; T2R14 agonist) in 16HBE cells with different CFTR genotypes as indicated. Right, bar graph showing DAF-FM change after 5 min stimulation with FFA. F508del and G542X CFTR cells exhibited lower NO responses compared with cells containing Wt CFTR; *p<0.05 by one-way ANOVA with Bonferroni posttest. Bar graph shows data points from n = 5 independent experiments using cells at different passages. **g** Left, DAF-FM trace of 16HBE cells during stimulation with 1 µg/ml or 10 µg/ml SC79. NO donor SNAP was used at the end of the experiment as a control. Right, bar graph showing DAF-FM fluorescence after 10 min stimulation with various concentrations of SC79; *p<0.05 compared with isogenic cells in the absence of SC79 (0 µg/ml) by one-way ANOVA with Bonferroni posttest. No differences were observed between genotypes at the same SC79 concentration by one-way ANOVA. Bar graph shows data points from n = 4 independent experiments using cells at different passages.

To continue to characterize the SC79 downstream NO production, we used primary nasal epithelial cells isolated from residual surgical material and cultured at air-liquid interface (ALI), which induces formation of motile cilia and goblet cells (**Figure 2a**), mimicking the in vivo nasal epithelium better than submerged cultures or cell line models [55, 65]. We loaded human nasal ALIs with DAF-FM to track NO production. SC79 dose-dependently increased DAF-FM fluorescence over >10 min (**Figure 2b**). The DAF-FM fluorescence increases were blocked by NOS inhibitor L-NAME but not by inactive analogue D-NAME (**Figure 2c**). DAF-FM fluorescence changes were also blocked by NO scavenger cPTIO (**Figure 2c**). DAF-FM fluorescence changes were also blocked by siRNA directed against eNOS but not against the neuronal NOS (nNOS) isoform (**Figure 2d**). These data together confirm that SC79 stimulates NO production in primary nasal cells via eNOS.

**Fig. 2.**
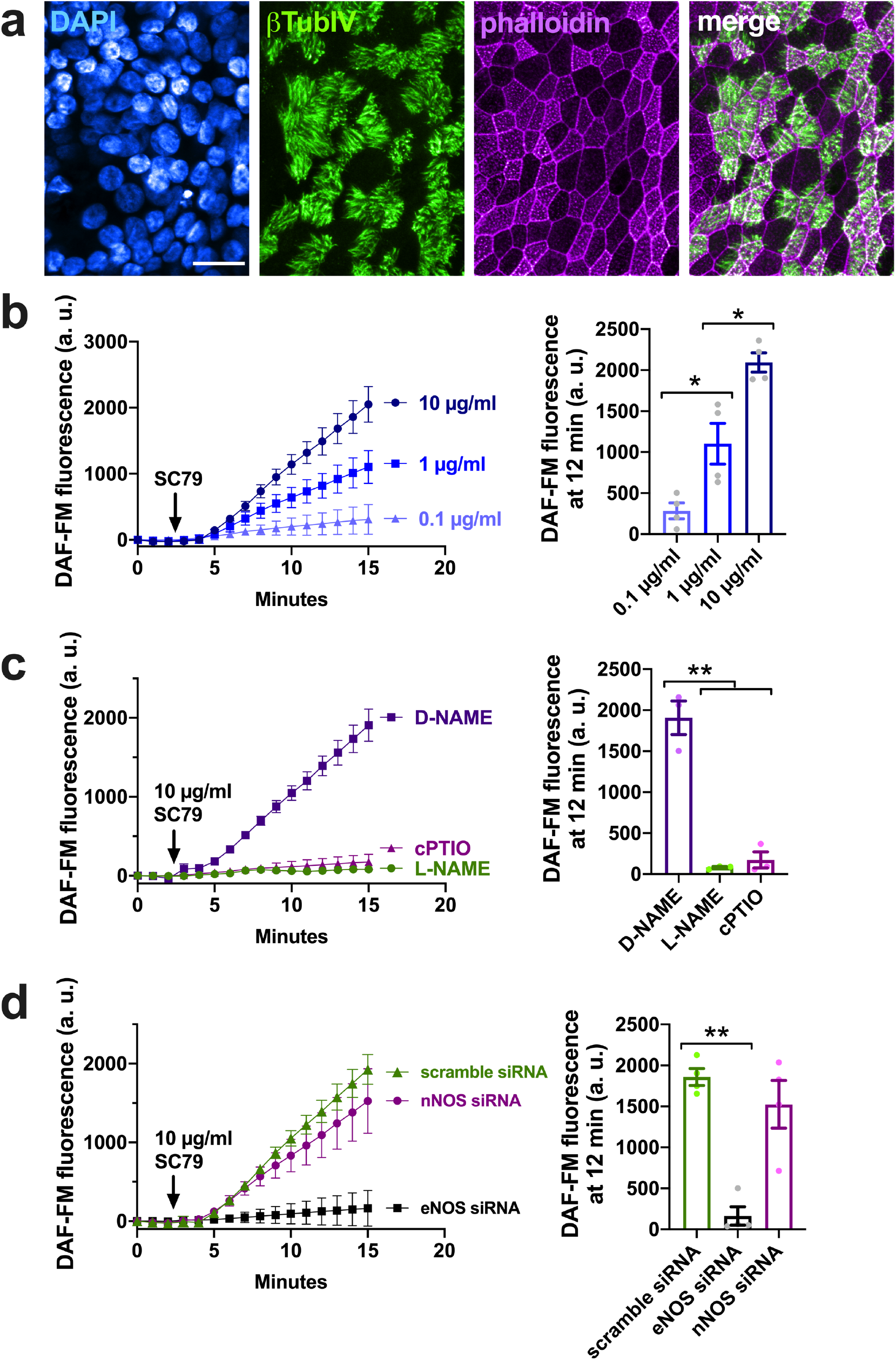
SC-79-induced NO production in primary nasal cells is dependent on eNOS activity. **a** Effects of SC79 were tested in air-liquid interface (ALI) cultures of primary nasal epithelial cells, which form tight junctions (labeled with actin stain phalloidin, shown in magenta) and differentiated ciliated cells (shown by immunofluorescence of cilia-enriched β-tubulin isoform IV) and non-ciliated (goblet cells). Scale bar is 20 µm. **b** Traces (left) and bar graph (right) of NO-sensitive dye DAF-FM in primary nasal ALIs confirming dose-dependent increase in DAF-FM fluorescence by SC79. SC79 activates Akt independent of other kinases (e.g., ERK) which may be activated during stimulation with receptor ligands such as EGF. Data from 4 independent expeirments using ALIs from different patients. Significance by one-way ANOVA with Bonferroni posttest; \**p*<0.05. **c** Traces (left) and bar graph (right) showing inhibition by NO scavenger cPTIO (10 µM) or NOS inhibitor L-NAME (45 min pre-treatment; 10 µM) but not inactive D-NAME (45 min pre-treatment; 10 µM); n = 4 independent experiments using ALIs from 4 separate patients. Significance by one-way ANOVA with Bonferroni posttest; \*\**p*<0.01. **d** Traces (left) and bar graph (right) showing inhibition by eNOS but not scramble or nNOS siRNA (treatment as described [89]); n = 4 independent experiments using ALIs from 4 separate patients. Significance by one-way ANOVA with Bonferroni posttest; \*\**p*<0.01.

We tested if the SC79-activated NO production required Akt. We found that SC79-stimulated DAF-FM fluorescence increases were inhibited by Akt inhibitors GSK690693 and MK2206 (**Figure 3a**). DAF-FM fluorescence increases were also blocked by LY294002 (**Figure 3b**), and inhibitor of PI3 kinase (PI3K) upstream of Akt. We previously found that SC79 induced Akt phosphorylation requires PI3K [45]. The SC79-induced DAF-FM fluorescence increases were also blocked by Akt inhibitor API-2 but not protein kinase C inhibitor Gö6983 (**Figure 3b**). DAF-FM fluorescence increases were also not blocked by phospholipase C inhibitor U73122 or its inactive analogue U73343 (**Figure 3c**). Together, these data suggest that SC79-stimulated eNOS activation is due to Akt and not activation of another kinase.

**Fig. 3.**
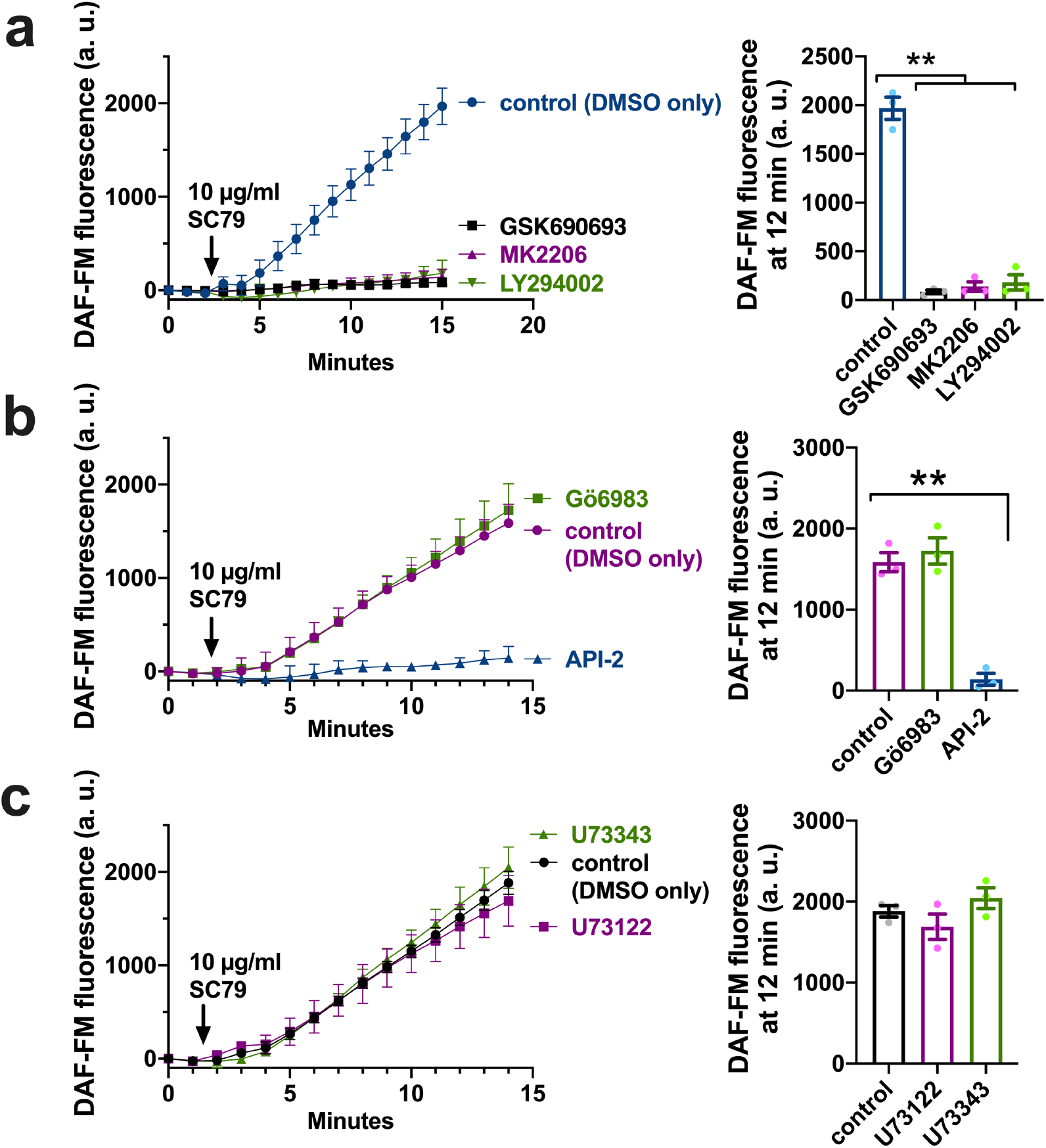
SC79 activation of eNOS requires Akt activity. **a** Trace (left) and bar graph (right) of DAF-FM fluorescence during stimulation with 10 µg/ml SC79 ± Akt inhibitor GSK6906932 (1 µM), Akt inhibitor MK2206 (1 µM), or LY294002 (1 µM). **b** Trace (left) and bar graph (right) of DAF-FM fluorescence during stimulation with 10 µg/ml SC79 in the presence of PKC inhibitor Gö6983 (1 µM) or Akt inhibitor API-2 (1 µM). **c** Trace (left) and bar graph (right) of DAF-FM fluorescence during stimluation with 10 µg/ml SC79 in the presence of PLC inhibitor U73122 (10 µM) or inactive analogue U73343 (10 µM). DMSO only was included in all control conditions. Pre-treatment with inhibitors was for 15 min prior to stimulation in the continued presence of inhibitor. All results from 3 independent experiments on different days using ALI from 3 different patients. Significance by one way ANOVA with Dunnett’s posttest comparing all values to control.

We previously showed that SC79 had no toxic effects on cell metabolism or proliferation at 24 and 48 hours [45]. To further test if acute SC79 stimulation had any toxic or pro-inflammatory effects on primary nasal cells, we treated ALIs with SC79 for 24 hours and saw neither LDH release (via colormetric assay; **Figure 4a**) nor IL-8 transcript production by qPCR (**Figure 4b**). We looked at secretion of several airway epithelial-derived (IL-8, IL-6, G-CSF, and GM-CSF [25, 66–72]) using ELISA. We saw no cytokine release with SC79. TLR5 agonist flagellin (derived from *Pseudomonas aeruginosa*) was used as a positive control. This agrees with our previous study showing that SC79 has anti-inflammatory effects through activation of Nrf-2 [45].

**Fig. 4.**
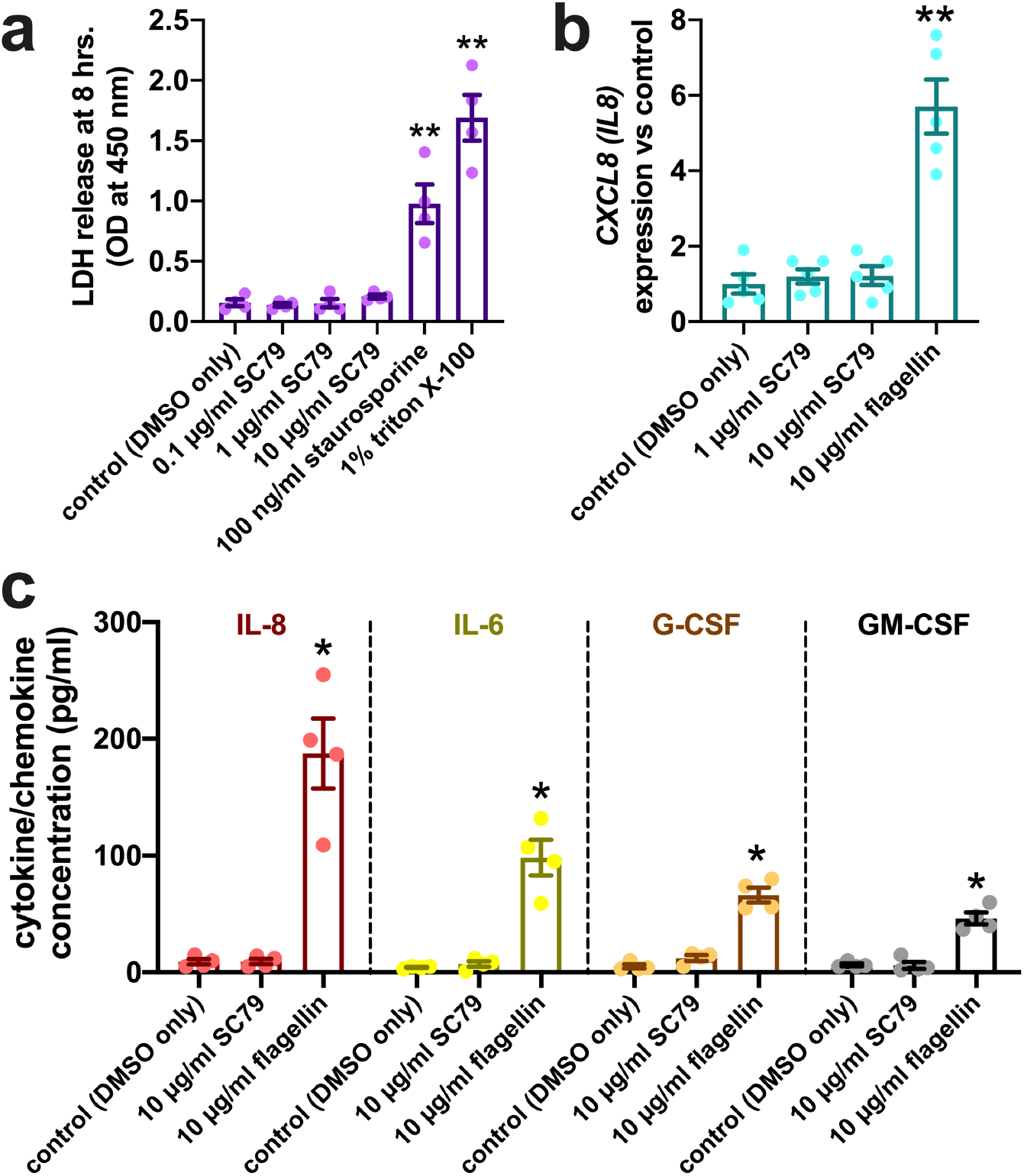
Lack of cytotoxic or inflammatory effects with acute exposure to SC79. **a** LDH release into cell culture media was measured via colorimetric assay. Staurosporine and triton X-100 were used as controls to induce apoptotic death (which is followed by secondary necrosis in the absence of phagocytic macrophages[90] and nonspecific lysis, respectively. No LDH release was observed with SC79 by one-way ANOVA; n = 4 ALIs per condition, each from a separate individual patient. **b** IL-8 transcription was quantified by qPCR (normalized to GAPDH) after 24 hours stimulation with SC79 or TLR5 agonist flagellin. Each data point is an independent ALI from a different patient (n = 5 total ALIs per condition). Significance by 1-way ANOVA with Dunnett posttest; \*\**p*<0.01. **c** Several epithelial-derived cytokines and chemokines (IL-8, IL-6, G-CSF, and GM-CSF) were tested with SC79 stimulation. No increases were observed with SC79 but all increased with flagellin; n = 4 ALIs per condition, each from a separate patient. Significance by one-way ANOVA with Bonferroni posttest.

With this lack of overt toxic effects with acute treatment, we hypothesized that short-term stimulation with SC79 might be useful to activate bactericidal NO production by nasal epithelial cells. To test if the NO produced during SC79 treatment entered the airway surface liquid (ASL), we used an assay to measure ASL NO using cell-impermeant RNS-sensitive dye DAF-2 overlayed in a small volume on top of the ALI cultures [22]. ASL DAF-2 fluorescence increased dose-dependently with SC79, and this was inhibited by NOS inhibitor L-NAME (**Figure 5a**). We also saw that ASL NO was not different in *TAS2R38* PAV/PAV vs AVI/AVI cultures, confirming that SC79 effects were independent of *TAS2R38* status and likely independent of T2R signaling (**Figure 5b**). We also saw no difference between ASL DAF-2 responses in non-CF vs CF (F508del/F508del) cultures (**Figure 5c**).

**Fig 5.**
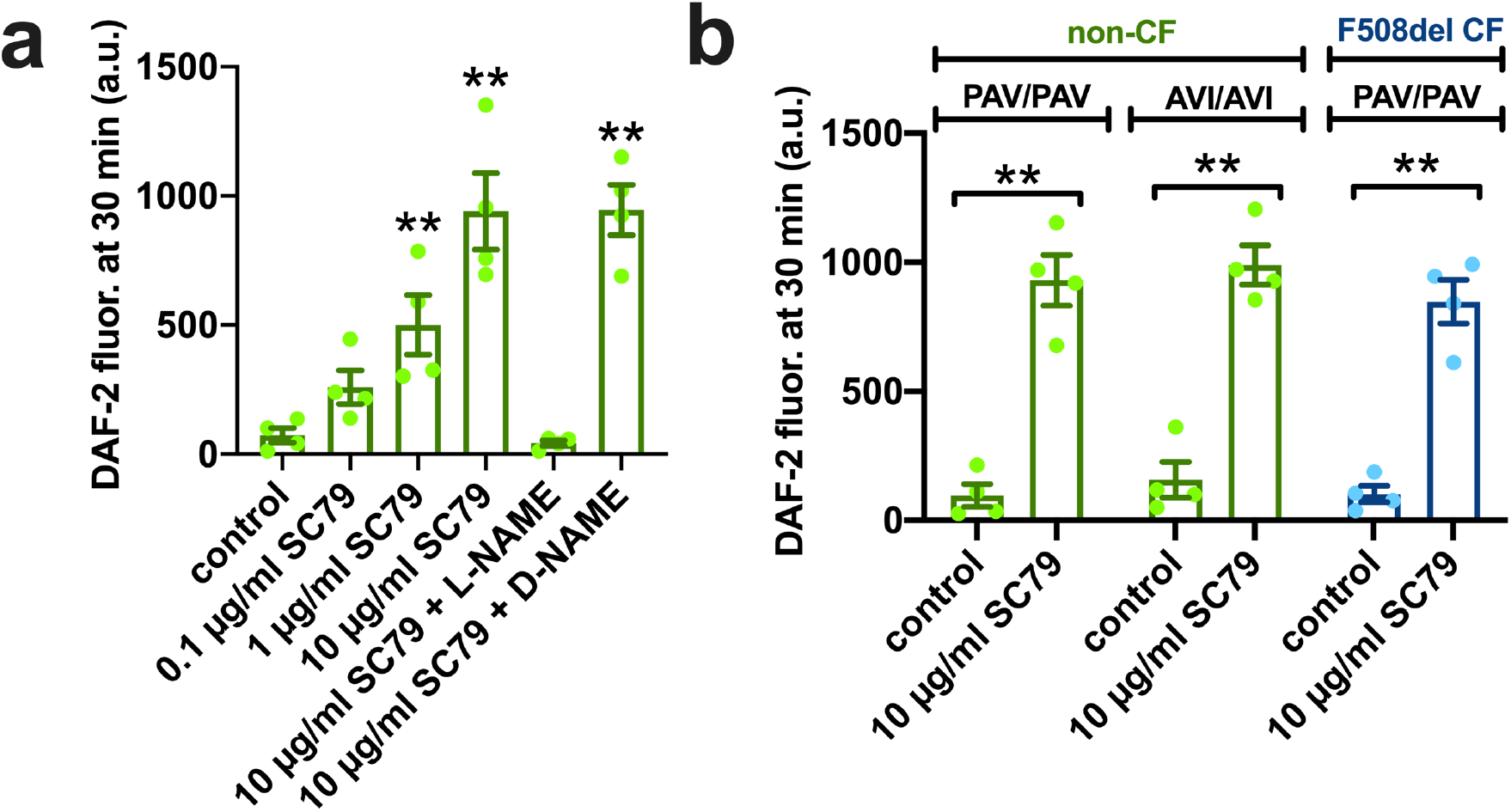
SC79 stimulates release of NO into the airway surface liquid. **a** Bar graph of fluorescence changes of cell impermeant DAF-2 overlayed on top of ALI cultures (as described [22]) showing dose-dependent increases with SC79 and inhibition by NOS inhibitor L-NAME but not inactive analogue D-NAME. Significance by one-way ANOVA comparing all values with control; **p<0.01; n = 4 ALIs per condition, each from a separate individual patient. **b** DAF-2 measurement of ASL NO production was tested in *TAS2R38* PAV/PAV (homozygous functional) vs AVI/AVI (homozygous non-functional) cultures. SC79 significantly increased DAF-2 fluorescence vs control in both genotypes; no difference was observed between SC79 stimulated PAV/PAV or AVI/AVI cultures. No difference was observed between non-CF and CF PAV/PAV cultures; n = 4 ALIs per condition, each from a separate individual patient. Significance by one-way ANOVA with Bonferroni posttest; **p<0.01.

Is this NO bactericidal? To test this, we used a bacterial killing assay in which nasal ALIs kill *P. aeruginosa* in an NO-dependent fashion [22, 25] with *TAS2R38* AVI/AVI (non-functional) ALIs, which do not respond to endogenous bacterial acyl-homoserine lactones or quinolones that activate T2R38. With no exogenous stimulation, AVI/AVI exhibit minimal bacterial killing in this assay [25]. Bacteria were incubated with nasal ALIs for 2 hours and collected and stained for live (Syto9) and dead (propidium iodide [PI]) bacteria. Lab *P. aeruginosa* strains PAO-1 and ATCC27853 were used along with clinical CRS isolates P11006, 2338, and L3847. SC79 stimulated dose-dependent killing (decreased Syto9/PI fluorescence ratio) of PAO-1 and ATCC27853 in an L-NAME-sensitive and MK2206-sensitive manner (**Figure 6a**). SC-79 also stimulated killing of clinical isolates (**Figure 6a**). Antibacterial effects were confirmed by counting CFUs, and less *P. aeruginosa* were recovered from AVI/AVI nasal ALIs treated with SC79 vs ALIs treated with vehicle (0.1% DMSO) only (**Figure 6b-c**). Overall, these results suggest SC79 can stimulate nasal epithelial cells to produce bactericidal levels of NO *in vitro*. We tested these antibacterial responses in ALIs derived from non-CF and CF (F508del/F508del) patients. We found that CF ALIs were equivalent to non-CF ALIs in terms of *P. aeruginosa* killing during SC79 stimulation (**Figure 6d**).

**Fig 6.**
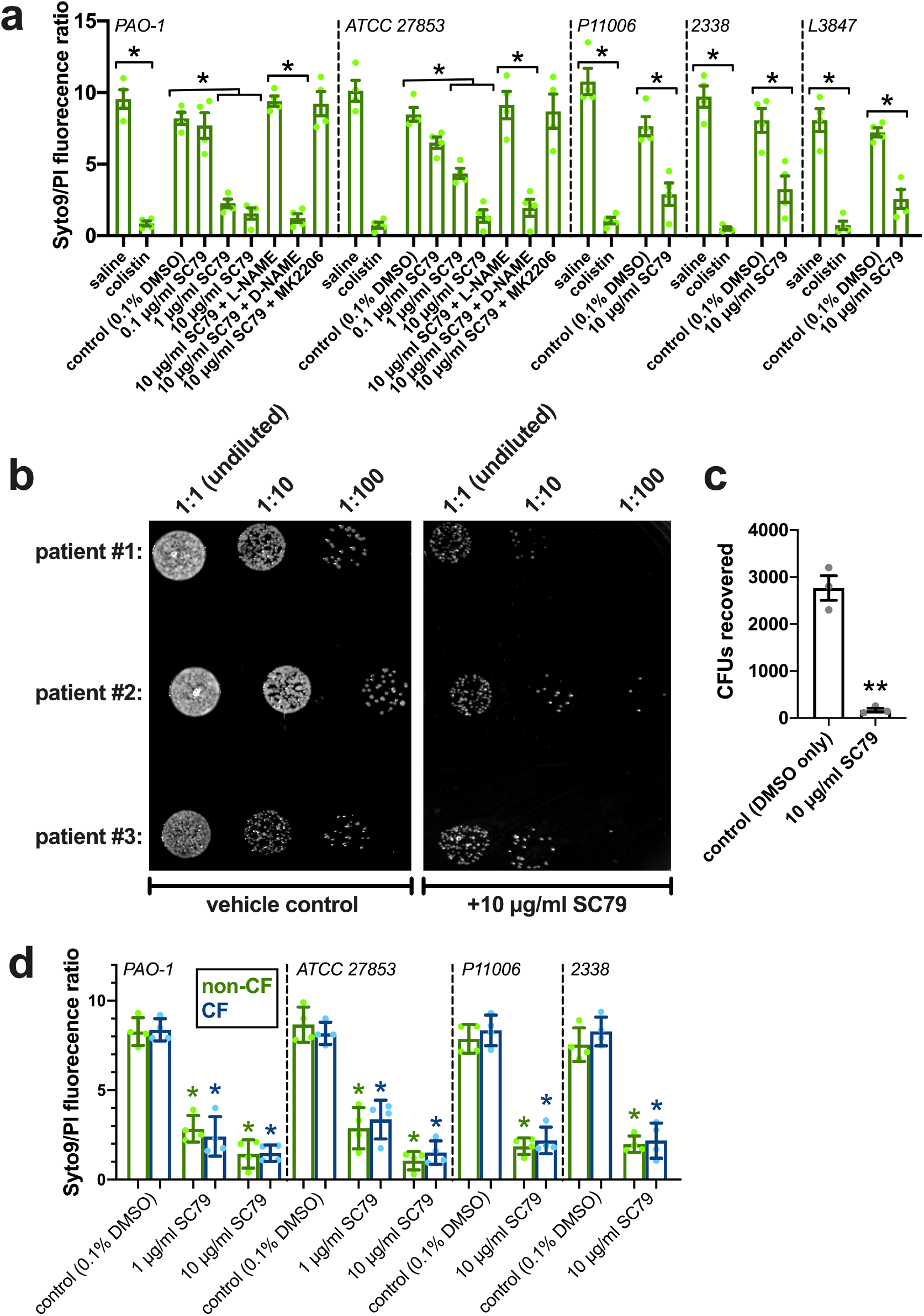
SC79-activated NO production has antimicrobial effects. **a** Bar graph of Syto9/PI fluorescence from bacterial killing assay (as described [22, 25]) using AVI/AVI *TAS2R38* cultures showing less viable bacteria (lower Syto9/PI ratio) with SC79. Significances by one-way ANOVA with Bonferroni posttest.All results in *A-C* from 4 independent experiments on different days using ALIs from 4 different cultures. **b** Bacteria recovered from nasal epithelial cultures (*TAS2R38* AVI/AVI) were collected, diluted in saline, and spotted on LB plates for manual CFU counting. **c** Quantification revealed a reduction in CFUs with SC79 treatment. Significance by Student’s *t* test; \*\**p*<0.01.

To further determine if beneficial immune responses might be observed, we tested if SC79 enhanced macrophage phagocytosis. Macrophages are important players in early airway innate immunity [73, 74]. T2R bitter taste receptors expressed in macrophages can also stimulate phagocytosis via NO signaling [31]. We cultured and differentiated human macrophages from healthy apheresis donors and measured NO production by DAF-FM fluorescence after 20 min stimulation with SC79. SC79 increased DAF-FM fluorescence in a MK2206-dependent, GSK690693-dependent, and L-NAME-sensitive manner (**Figure 7a**).

**Fig 7.**
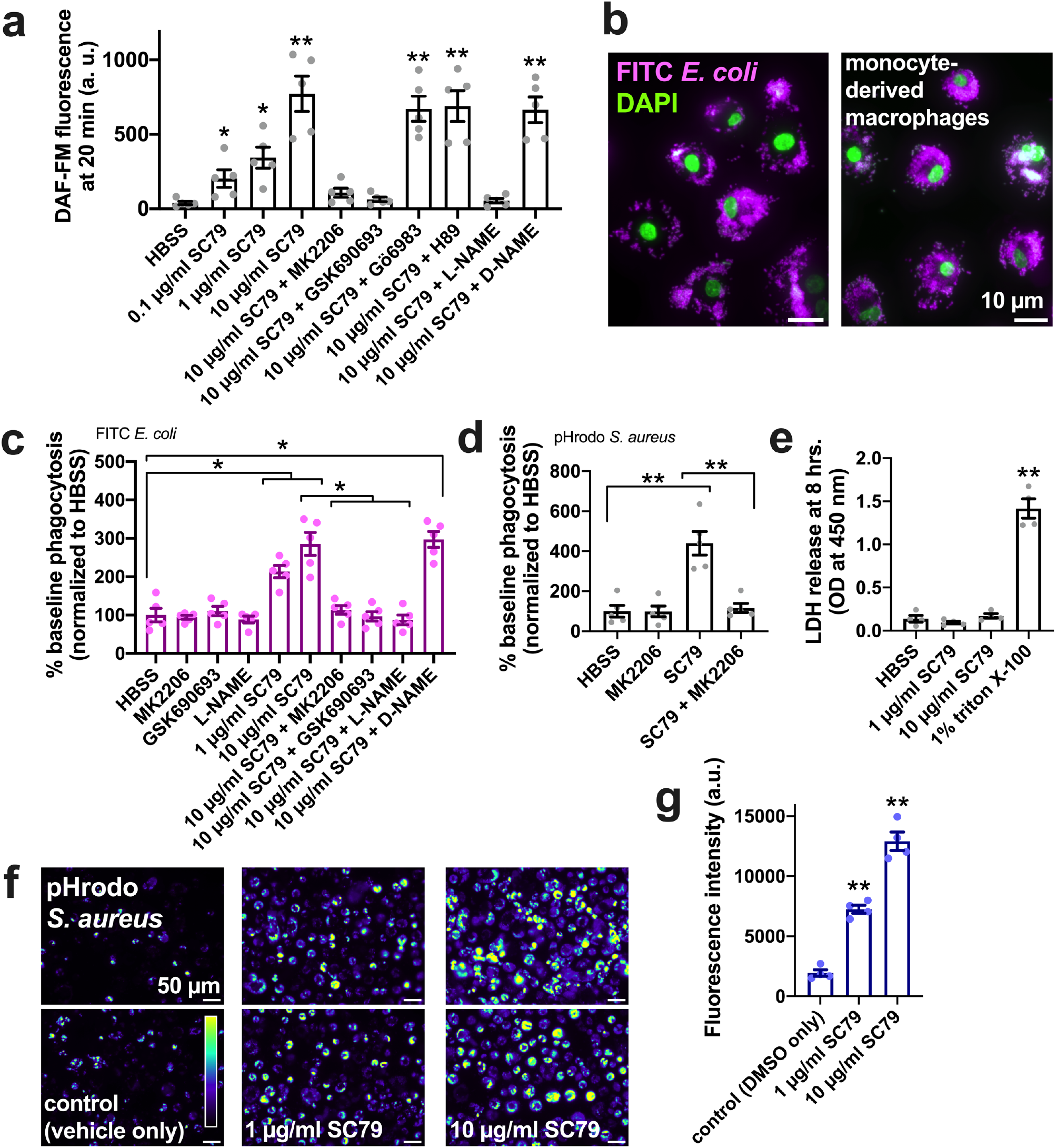
SC79 stimulates macrophage phagocytosis via Akt and NO signaling. **a** Bar graph showing DAF-FM fluorescence measured by microscopy 30 min after stimulation with compounds as indicated (MK2206, GSK690693, L-NAME, D-NAME, and Gö6983 used as previously described in the study). Gö6983 (PKC inhibitor) and H89 (1 µM; PKA inhibitor) used as controls. **b** Images showing phagocytosis of FITC-labeled *E. coli* (magenta, DAPI nuclear stain in green) in primary human monocyte-derived macrophages, as described.[22, 31] **c** FITC fluorescence (indicating macrophage phagocytosis) increased with SC79 treatment, which was blocked by Akt inhibitors MK2206 or GSK690693 and NOS inhibitor L-NAME. Significance by one-way ANOVA with Dunnett’s posttest comparing all values to control; \**p*<0.05. Data are from 5 independent experiments using cells from 5 donors. **d** Phagocytosis of pHrodo S. aureus also increased with 10 µg/ml SC79 and was blocked by MK2206. Note that pHrodo only fluoresces in acid environments like the phagosome, confirming that internalization reflects phagocytosis. Data from 5 independent experiments using cells from 5 donors. Significance by one-way ANOVA with Bonferroni posttest; \*\**p*<0.01. **e** LDH release from macrophages was measured after 8 hours stimulation. Data from 4 independent experiments using cells from 4 donors. Significance by one-way ANOVA with Dunnett’s posttest comparing all values to control; \*\**p*<0.01. **f** Microscope images of pHrodo *S. aureus* phagocytosed in macrophages. **g** Quantification of 4 independent experiments as shown in *E* confirms dose-dependent increase in phagocytosis with SC79. Significance by one-way ANOVA with Dunnett’s posttest comparing all values to control; \*\**p*<0.01.

We next measured phagocytosis of FITC-labled *Escherichia coli* bioparticles (**Figure 7b**) with fluorescence read on a plate reader. SC79 increased macrophage phagocytosis in a manner inhibited by AKT inhibitors MK2206 and GSK690693 as well as NOS inhibitor L-NAME (**Figure 7c**). SC79 stimulated macrophage phagocytosis was also confirmed by a plate reader assay of *Staphylococcus aureus* labeled with pH-sensitive dye pHrodo (**Figure 7d**), suggesting increased phagocytosis with SC79 and inhibition by MK2206. No LDH release was observed from macrophages over 8 hours stimulation with SC79, suggesting no overt toxic effects (**Figure 7e**). SC79-stimulation of phagocytosis of *S. aureus* was also confirmed by direct microscopy visualization and quantification, showing dose dependent effects of SC79 at 1 and 10 µg/ml (**Figure 7f-g**).

## Discussion

The nose is the “front line” of respiratory defense, where host-pathogen interactions occur with every breath [75]. Failure of upper respiratory defenses can lead to microbial overload and chronic rhinosinusitis (CRS), a complex syndrome of sinonasal infection and/or inflammation affecting >35 million Americans. CRS creates an aggregated direct healthcare cost of ∼$8 billion annually [76], with indirect costs (e.g., missed workdays and decreased productivity) adding to the >$20 billion socioeconomic burden [76]. Importantly, conventional management of acute and chronic rhinosinusitis accounts for nearly 20% of adult antibiotic prescriptions [76], making CRS a major contributor to the antibiotic resistance crisis, called “arguably the greatest risk…to human health” by the World Economic Forum [77]. As the prevalence of resistance increases, conventional CRS therapies are becoming less effective [44, 78, 79]. Stimulating endogenous host defenses is an attractive alternative strategy to boost immunity without the pressures for single-agent resistance. NO is an important component of nasal innate defense, and our data suggest that activation of Akt via a small molecule like SC79 may have acute antibacterial effects in the nasal epithelium by activating NO production.

While it is likely that one or more of the three Akt isoforms play an important role in every cell type in the body, Akt signaling is understudied in the context of airway innate immunity or airway epithelial physiology in general. The role of Akt in innate defense is unclear, even despite Akt being downstream of several toll-like receptors [64, 80]. The study of Akt’s role is complicated by cross talk between multiple convergent signaling pathways that are activated downstream of TLRs and other receptors. We previously examined Akt signaling in airway cells using SC79 to directly activate Akt in the absence of other kinases [45], and interestingly found anti-inflammatory effects of SC79 Akt activation due to Nrf2 upregulation and translocation to the nucleus [45]. Here, we show that the NO produced during SC79 stimulation of primary nasal ALIs is Akt-dependent and sufficient to kill bacteria. It may also be sufficient to inactivate viruses or fungi, but further testing is needed. SC79 also enhanced macrophage phagocytosis via an Akt-dependent and NO-dependent mechanism. Thurs, we hypothesize that SC79 delivered topically in a nasal spray or rinse might be effective against some types of upper respiratory infections by stimulating endogenous NO production.

Notably, Akt plays a central role in cell growth and metabolism.[47] An important caveat to activating Akt hinges on the central role that the PI3K-Akt pathway plays in driving tumorogenesis or tumor proliferation in many cancers [81, 82]. Activating mutations upstream of Akt in receptors like EGFR or downstream PI3K are frequently found in cancers, making these proteins important targets to inhibit in many cancers [81]. However, we previously did not observe increased cellular proliferation with SC79 during acute (24-48 hrs) stimulation [45], and to our knowledge tumors have not been reported in animal models in which SC79 has been administered [58–61]. We hypothesize that transient pharmacological Akt activation in an acute infection setting will likely have very different effects compared with activating mutations that cause chronic Akt activation and which are often accompanied by other oncogenic driver mutations in parallel. However, the use of SC79 still requires further *in vitro* and *in vivo* testing. However, our data here and from a previous study [45] suggest acute SC79 may have both some anti-inflammatory and some bactericidal benefits with limited cellular toxicity in a topical delivery setting for respiratory infections.

## Conclusions

While NO donors are being actively explored for CRS [83, 84], we hypothesize that a useful alternative strategy is to target endogenous signaling pathways to generate more physiological levels of NO production. An example of this is the cilia T2R-dependent pathway [28]. When this pathway has reduced function (in *TAS2R38* AVI/AVI patients), several measures of CRS disease severity are worsened [32, 39, 85–87]. One strategy to overcome this is to use T2R agonists like quinine [88] or diphenhydramine derivatives [56] that might activate T2Rs in cilia beyond T2R38. Still another strategy, explored here, is using a compound like SC79 that signals to eNOS via a completely different mechanism (Akt signaling) from T2Rs (Ca^2+^/calmodulin signaling). We believe that the above results comprise pilot data suggesting Akt activation should be further explored as a non-antibiotic antimicrobial strategy in the context of respiratory infections in CRS and possibly other airway diseases.

## Acknowledgements

This study was supported by NIH grants R01DC016309 (to R.J.L.), R01AI167971 (to N.D.A., J.N.P., and R.J.L.), and Cystic Fibrosis Foundation grant LEE21G0 (to R.J.L.). We thank J. Riley (University of Pennsylvania Perelman School of Medicine, Human Immunology Core, supported by P30CA016520 and P30AI045008) for primary human monocytes. We thank M. Victoria and J. Freund (University of Pennsylvania Perelman School of Medicine) for technical assistance. We thank B. Chen, N. Cohen, and L.E. Kuek (University of Pennsylvania Perelman School of Medicine) for assistance with isolation and culture of patient nasal cells.

## CRediT authorship contributions

Robert J. Lee: Conceptualization, Investigation, Methodology, Formal analysis, Writing – original draft, Writing – review & editing, Project administration, Funding acquisition. Nithin D. Adappa: Resources, Data curation, Project administration, Writing – review & editing, Funding acquisition. James N. Palmer: Resources, Data curation, Project administration, Writing – review & editing, Funding acquisition.

## Availability of data and materials

All data generated or analyzed during this study are included in this article and its additional files.

## Declarations

### Ethics approval and consent to participate

Written informed consent was obtained from patients undergoing medically necessary sinonasal surgery in accordance with the U.S. Department of Health and Human Services Title 45 CFR 46.116, the Declaration of Helsinki, and the University of Pennsylvania guidelines regarding residual clinical material in research under IRB protocol #800614 **Consent for publication:** Not applicable.

### Availability of data and materials

All data generated or analyzed during this study are included in this article and its supplementary files.

### Corresponding author(s)

Robert J Lee, Department of Otorhinolaryngology, Hospital of the University of Pennsylvania, 3400 Spruce Street, 5^th^ floor Ravdin Suite A, Philadelphia, PA USA 19104. Phone 215-573-9766.

